# Consistent local dynamics in the brain across sessions are revealed by whole brain modeling of resting state activity

**DOI:** 10.1101/104232

**Authors:** Patricio Donnelly Kehoe, Victor M. Saenger, Nina Lisofsky, Simone Kühn, Morten L. Kringelbach, Jens Schwarzbach, Gustavo Deco

## Abstract

Resting state fMRI has been the primary tool for studying the functional organization of the human brain. However, even at so-called “rest”, ongoing brain activity and its underlying physiological organization is highly dynamic and yet most of the information generated so far comes from group analysis. Here we developed an imaging-based technique capable of portraying information of local dynamics at a single-subject level reliably by using a whole-brain model that estimates a local bifurcation parameter, which reflects if a brain region presents stable, asynchronous or transitory oscillations. Using 50 longitudinal resting state sessions of one single subject and single resting state sessions from a group of 50 participants we demonstrated that individual global and local brain dynamics can be estimated consistently with respect to a reference group using only a scanning time of 15 to 20 minutes. We also showed that brain hubs are closer to a transition point between synchronous and asynchronous oscillatory dynamics and that dynamics in frontal areas have larger variations compared to other regions. Finally, we analyzed the variability and error of these dynamics and found high symmetry between hemispheres, which interestingly was reduced by adding more sessions. The framework presented here can be used to study functional brain dynamics on an individual level, opening new avenues for possible clinical applications.

**Bullet points:** Local brain dynamics are consistent across scans.

Four scans of five minutes each are enough to get highly reliable and consistent results.

Hub areas are in a transition point between a synchronous and asynchronous regime.

Variability and error of local dynamics presented high symmetry between hemispheres.

## 1 Introduction

Resting state functional MRI (RS-fMRI) enables the detailed description of group-level functional brain organization at multiple spatial scales (Deco et. al., 2011; Singer et. al., 2013). Additionally, at the group level, many studies have addressed the intrinsic functional architecture of the healthy brain (e.g. Damoiseaux et. al., 2006; Honey et. al., 2010) while others have found significant differences between control participants and patients suffering from a wide variety of brain disorders, e.g. schizophrenia (Lynall et. al., 2010; Damaraju et. al., 2014), Parkinson’s Disease (Göttlich et. al., 2013) and autism (Jones et. al., 2010). Although these studies show that group-level analyses provide important information about the overall organization of the brain, few of them have applied a method at a single-subject level with high consistency, and in a way that can be used for clinical purposes (but see Mueller et. al., 2013; Laumann et. al., 2015).

This represents a major issue given the well-known large anatomical and functional variability across individuals (Frost et. al., 2012). Moreover, functional variability is also expected within a single subject depending on the state of the subject and on environmental noise (Mueller et. al., 2013). A reliable single-subject analysis of brain dynamics should then yield stable and reproducible results across different scan sessions (Fiecas et. al., 2013; Zuo et. al., 2014), while results also need to be consistent enough within a given group (Castellanos et. al., 2013; Ferreira et. al., 2013). Previous studies that analyzed the functional connectivity (FC) of the brain using a correlation matrix (CM) or using more advanced statistical techniques such as Independent Component Analysis (ICA) (Damoiseaux et. al., 2006; reviewed in Fox and Raichle, 2007) hypothesized that functional connectivity is constant over time, although it is now becoming clear that even at rest, there are dynamical processes worth considering (Hutchison et. al., 2013). Conversely, numerous studies have postulated that brain dynamics are essential to explain both health and disease, placing phase synchronization fluctuations as one of the main features required to understand how integration and segregation arise and modulate brain states (Tognoli & Kelso, 2014). For this reason, some groups have proposed that functional connectivity dynamics (FCD) have to be taken into account to develop FC-related metrics that properly reflect the dynamical nature of brain function (Calhoun et. al., 2014; Deco et. al., 2014; Deco et. al., 2015; Hansen et. al., 2015).

In this context, whole-brain computational models have been used as powerful tools to understand the relation between structural and functional brain connectivity by linking brain function with its physiological underpinnings (Ponce-Alvarez et. al., 2015; Deco et. al., 2015; Kringelbach et. al., 2015). In contrast to conventional resting state analyses, studies based on brain models allow the exploration of brain dynamics through parameter tuning (Deco et. al., 2014; Deco et al., 2016) both in health and disease (Cabral et. al., 2013). A recent study (Deco et al., 2016) used a novel whole-brain modeling technique to simulate brain dynamics estimating a local parameter which contains information about the oscillatory nature of a given region. In this work we used this model and further developed a processing method to provide new insights about brain function at a single-subject level as well as to estimate consistency of local dynamics to explore the minimum scan length and number of sessions required to have a robust estimation. We showed that these dynamics are key features for both global and local consistency exploration and that longer scanning times might reveal important information impossible to uncover with single-session analysis or short scans.

### 1.1 Conceptual considerations of the whole-brain Hopf bifurcation model

In order to acquire flexible and consistent results through scanning sessions, we developed and applied a model (Deco et al., 2016) that uses functional and structural connectivity information to simulate whole brain activity. As shown in Figure 1, the model uses brain dynamics from fMRI data and the structural connectivity from diffusion weighted imaging (DWI) data to construct an interconnected network (Figure 1.A). In this network, each node (brain region) presents supercritical Hopf bifurcation dynamics (Marsden et. al., 2012) that depending on the value of its bifurcation parameter (BP) reflects if the region behaves either in a noisy asynchronous or stable oscillatory manner (Figure 1.B).

The model was fitted to the empirical data (BOLD signal) using two parameters. The first is the *global scaling factor (G)*, that determines the impact of the structural connectivity to brain dynamics. The other is a set of local (i.e. per-node) bifurcation parameters (BP) constituting the brain’s Dynamical Core (DynCore). Figure 1B illustrates the notion that a negative value of a BP means that this particular node is working as a noise generator, while a node with positive BP generates stable oscillatory dynamics. Lastly, nodes that exhibit a BP around zero can be understood as flexible systems on the edge of a bifurcation capable of jumping from one state to the other depending on physiological demands. An in-depth mathematical explanation of the Hopf-bifurcation model can be found in the methods section. Our model allows one to study the brain locally (at a region scale), but also globally using the whole DynCore. A “flexible brain” at rest is expected to have a global mean of the bifurcation parameters around zero in order to rapidly adapt to environmental or internal demands.

**Figure 1:**
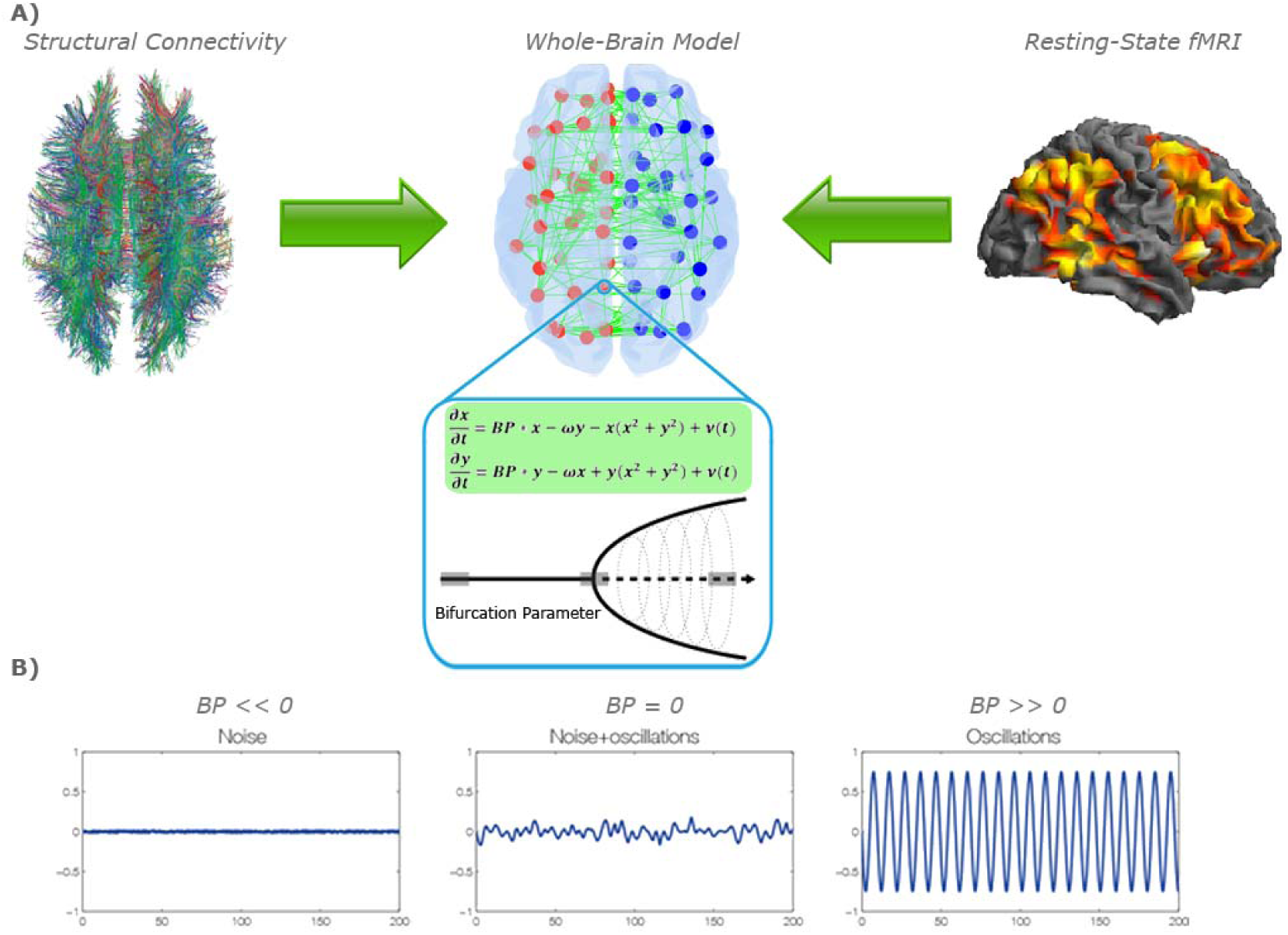
*Whole-brain model*. A) The whole-brain model was based on the structural connectivity (SC) matrix derived from tractography of DWI (left) between the 90 regions of the Automated Anatomical Labeling (AAL) parcellation (middle). The control parameters of the models were tuned using the grand average FC and FCD derived from fMRI BOLD data (right). B) For modelling local neural masses we used the normal form of a Hopf bifurcation, where depending on the bifurcation parameter (BP in the equations – green box–), the local model generates a noisy signal (left), a mixed noisy and oscillatory signal (middle) or an oscillatory signal (right). It is at the border between noisy and oscillatory behavior (middle), where the simulated signal looks like the empirical data, i.e. like noise with an oscillatory component around 0.05 Hz.

To allow comparing dynamics across sessions and participants we further developed a normalization method of the bifurcation parameters. We aimed to demonstrate that DynCores obtained using normalized bifurcation parameters (NBPs) provide a framework that can quantify brain dynamics consistent enough to be compared across participants and sessions. To test this hypothesis we applied the model to 50 longitudinal resting state (RS) sessions from a single subject and compared the within-subject consistency to that estimated using 50 recordings from different participants. Finally, we searched for an optimal number of sessions required for accurately estimate parameters, which could ultimately be used as local estimators of the brain dynamics with high test-retest reliability.

## 2 Materials and methods

### 2.1 Participants

Fifty one participants were recruited in total from which one participant volunteered to be included in the longitudinal part of the study in which she was scanned 50 times over the course of 6 months (female, aged 29). The remaining fifty participants (all female, mean age 24, SD=3.14, range: 18−32) underwent scanning with the same MRI sequences only once.

The study was approved by the local ethics committee (Charité University Clinic, Berlin). Participants gave written consent. The other participants were asked on the phone during recruitment whether they ever had a psychiatric disease and negated that. Other medical and neurological disorders were also reasons for exclusion. None of the participants showed structural abnormalities in the MRI scans.

### 2.2 Scanning Procedure

Images were collected on a 3T Verio MRI scanner system (Siemens Medical Systems, Erlangen, Germany) using a 12-channel radiofrequency head coil. First, high-resolution anatomical images were acquired using a three-dimensional T1-weighted magnetization prepared gradient-echo sequence (MPRAGE) with a repetition time = 2300 ms, an echo time = 3.03 ms, flip angle = 9°, 256 × 256 × 192 matrix and a 1 × 1 × 1 mm voxel size. Whole brain functional images were collected using a T2*-weighted EPI sequence sensitive to BOLD contrast (TR = 2000ms, TE = 30ms, image matrix = 64 × 64, FOV = 224 mm, flip angle = 80º, slice thickness = 3.5 mm, 35 near-axial slices, aligned with the AC/PC line, 150 volumes, 5 min duration).

#### 2.2.2 DWI data used on the model

The structural connectivity data was obtained from participants from a different study recruited at Aarhus University. Data were collected at the Center of Functionally Integrative Neuroscience (CFIN), Aarhus University, Denmark, from 16 healthy right-handed participants (11 men and 5 women, mean age: 24.75, SD=2.54). Participants with psychiatric or neurological disorders (or a history thereof) were excluded.

The structural connectivity data (structural MRI and DWI) were collected on a 3T Siemens Skyra scanner system (Siemens Medical Systems, Erlangen, Germany). The structural MRI T1 parameters: voxel size of 1 mm^3^; reconstructed matrix size 256x256; echo time (TE) of 3.8 ms and repetition time (TR) of 2300 ms. The DWI data were collected using TR = 9000 ms, TE = 84 ms, flip angle = 90°, reconstructed matrix size of 106x106, voxel size of 1.98x1.98 mm with slice thickness of 2 mm and a bandwidth of 1745 Hz/Px. Furthermore, the data were collected with 62 optimal nonlinear diffusion gradient directions at *b*=1500 s/mm^2^. One non-diffusion weighted image (*b=*0) per 10 diffusion weighted images was acquired. Additionally, the DWI images were collected with different phase encoding directions. One set was collected using anterior to posterior phase encoding direction and the second acquisition was performed in the opposite direction.

### 2.3 Dataset and preprocessing

#### 2.3.1 Empirical fMRI Data Analysis

The first ten volumes were discarded to allow the magnetization to approach a dynamic equilibrium. Data pre-processing, including slice timing, head motion correction (a least squares approach and a 6-parameter spatial transformation) and spatial normalization to the Montreal Neurological Institute (MNI) template (resampling voxel size of 3 mm × 3 mm × 3 mm), were conducted using the SPM5 and Data Processing Assistant for Resting-State fMRI (DPARSF, Chao-Gan and Yu-Feng, 2010). A spatial filter of 4 mm FWHM (full-width at half maximum) was used. One fMRI recording with head motion above 3 mm of maximal translation (in any direction of x, y or z) and 1.0° of maximal rotation throughout the course of scanning was excluded.

After pre-processing, linear trends were removed. Then the fMRI data was temporally band-pass filtered (0.01 - 0.25 Hz) to reduce low-frequency drift and high-frequency respiratory and cardiac noise (Biswal et. al., 1995). The data was parcellated into regions of interest (ROIs) using the Automated Anatomical Labeling (AAL) atlas (Tzourio-Mazoyer et. al., 2002). Each recording was represented by 90 nodes with 140 time points each node.

#### 2.3.2 Structural Connectivity Data

A single average structural connectivity (SC) matrix was used to obtain all DynCores. This average SC matrix was obtained using DWI data, which has been computed in the context of a different study (Deco et al, in review). Briefly, we generated the structural connectivity maps for each participant using the same AAL90 template regions used in the functional MRI data. We computed the connections between regions in the whole-brain network (i.e. edges) probabilistic tractography using FSL’s bedpostx and probtrackx (Behrens et. al., 2003; Behrens et. al., 2007) in both datasets acquired (each with different phase encoding to optimize signal in difficult regions). Finally, data was averaged across participants.

### 2.4 Whole-Brain Model

The whole-brain model consists of 90 coupled brain areas (nodes) derived from the parcellation explained above. The global dynamics of the whole-brain model used here results from the mutual interactions of local node dynamics coupled through the underlying empirical anatomical SC matrix *C*. Each element *C*_*ij*_ of the matrix *C* denotes the density of fibers between cortical area *i* and *j* as extracted from the DTI based tractography (scaled to a heuristic maximum value of 0.2). The local dynamics of each individual node is defined by the normal form of a supercritical Hopf bifurcation, which is able to describe the transition from asynchronous noisy behavior to full oscillations. Thus, in complex coordinates, each node *j* is described by Equation 1.

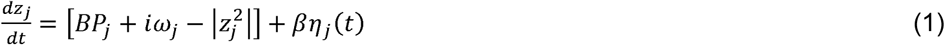

where 

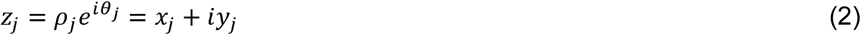

The term *η_j_*(*t*) is additive Gaussian noise with standard deviation *β*=0.02. *BP*_*j*_ is the node’s bifurcation parameter value, which reflects the local working point ultimately representing the region’s role in brain dynamics. Specifically, this normal form has a supercritical bifurcation at *BP*_*j*_ = 0, so that the local dynamics have a stable fixed point at *z*_*j*_ = 0 (which, because of the additive noise, corresponds to an asynchronous state) and for *BP*_*j*_ > 0 there exists a stable limit cycle oscillation with frequency 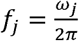. Thus, the whole-brain dynamics is defined by the set of Equations 3 and 4.

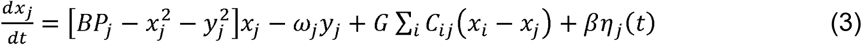

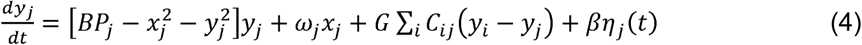

In the latter equations, *G* is a *global scaling factor* (global conductivity parameter scaling equally all synaptic connections), which quantifies the importance of the structural information in brain dynamics. A higher optimal *G* means that modeled FC is more constrained by the SC than a lower optimal *G* in which structural information becomes less relevant. This extrinsic global scaling factor and the bifurcation parameters represent the collection of control parameters with which we study the optimal dynamical working point where the simulations maximally fit the empirical FC and the FCD. Using this information, we were able to model the BOLD signal of each node *j* with the variable *x*_*j*_.

The empirical BOLD signals were band-pass filtered between 0.04–0.07 Hz. This frequency band has been mapped to the gray matter and it has been shown to be more reliable and functionally relevant than other frequency bands (Biswal, 1995; Achard, 2006; Buckner, 2009; Glerean, 2012). Using this filter, the intrinsic frequency *ω_j_* of each modeled signal in each node is in the 0.04-0.07Hz band (*j*=1,…, n). The amplitude of the intrinsic frequencies was estimated from the data, as given by the averaged peak frequency of the narrowband BOLD signals of each brain region.

#### 2.4.1 Grand average FC and FCD matrices and similarity scores

The grand average FC is defined as the matrix of correlations of the BOLD signals between a pair of brain areas over the whole time window of acquisition. We computed a measure of similarity between the modeled and empirical FC by using the Pearson correlation coefficient of the correlation values within the matrix (Deco & Kringelbach, 2014; Deco et. al., 2014). This similarity score, that we called *Fitting*, reflects the static grand-average similarity between two vectorized matrices in terms of a Pearson correlation coefficient.

In order to characterize the time dependent and dynamic functional structure and similarity of the resting fluctuations, we first estimated both, modeled and empirical, dynamical-FC matrices (FCD) based on the phase difference over time. We then assessed the instantaneous phase *φ* of each node at every time point *t* by applying a Hilbert transform. Then, by using a six-seconds sliding window technique, we reconstructed a phase difference matrix that we used as a way to measure phase synchronization across time, which yields a dynamical FC or FCD (Deco et al., 2016). To evaluate a similarity score for this dynamical property, we computed the Kolmogorov-Smirnov distance between the empirical and modeled FCD (*KSDist*). Both the grand average FC and the FCD matrices were estimated for each step of *G*.

#### 2.4.2 Metastability

For each time point we computed the global synchronization between areas by means of the Kuramoto order parameter. The fluctuation of that parameter quantified by its standard deviation in time represents the global *metastability*, which depicts the variability of phase synchronization between nodes across time (Wildie, 2012). The Kuramoto order parameter is defined by Equation 5.

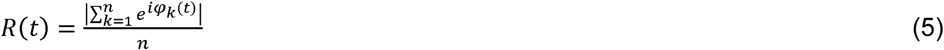

where *φ_k_*(*t*) is the instantaneous phase of each narrowband BOLD signal at node *k.* Under complete independence, the *n* phases are uniformly distributed and thus *R* is nearly zero, whereas *R* = 1 if all phases are equal (full synchronization).

#### 2.4.3 Local Optimization of Brain Nodes

The local optimization of each bifurcation parameter *BP*_*j*_ is based on the fitting of the spectral information from the empirical BOLD signals in each node. In particular, we aimed to fit the proportion of power in the 0.04-0.07 Hz band with respect to the 0.04-0.25 Hz band (i.e. we removed the smallest frequencies below 0.04 Hz and considered the whole spectra up until the Nyquist frequency, which is 0.25 Hz). For this, we filtered the BOLD signals in the 0.04-0.25 Hz band, and calculated the power spectrum *P*_*j*_(*f*) for each node *j*. This proportion is defined by,

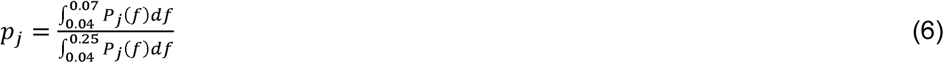

and updated the local bifurcation parameters by a gradient descent strategy, i.e.:

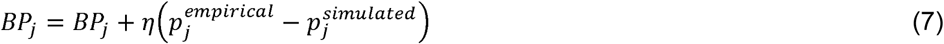

until convergence (a heuristic rule of 200 iterations). We used a fixed value of *η* = 0.005. For each G, we always considered the optimized set of BPs as the ones with the lower spectral distance *SpD*, where

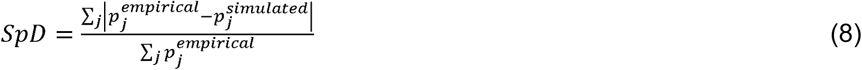

#### 2.4.4 Whole-brain model as a processing approach

Using cortical and subcortical nodes of one hemisphere at a time, we optimized the bifurcation parameters (BPs) and the simulated functional connectivity matrix for each of the empirical RS-fMRI recordings. To do this, we explored the coupling parameter space between 0 and 12 using steps of 0.1. Based on the previously described three similarity scores and having the considerations that i) metastability is desired to be higher (closer to 0.2) (Tognoli & Kelso, 2014), ii) fitting is better as it gets closer to 1 and iii) that the KSDist is better as it gets closer to 0 being the most important score as it portrays dynamics information, we developed a fourth score, the Global Similarity (*GS*) to express all these conditions in a single numerical value (Equation 9). In order to give KSDist more importance in the score we make *GS* depend on it quadratically, while it depends on the Fitting and the Metastability only linearly.

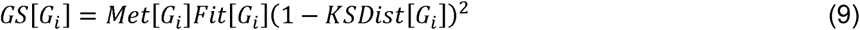

where *Met*[*G*_*i*_] and *Fit*[*G_i_*] represent the Metastability and the Fitting for a given *G_i_*. Then we used *GS*[*G*] to find the optimal *G*_*opt*_ in an automated way by difining it as the value of *G*_*i*_ where *GS*[*G*] exhibited its maximum value.

The output data from *G*_*opt*_ is a DynCore and a simulated functional CM. Results with a KSDist value larger than 0.3 or a Fitting smaller than 0.25 were discarded as we considered the RS recording to be noisy and unsuitable for the model to further generate acceptable simulations of brain dynamics. In this work, more than 4000 simulations were generated with a rejection rate of around 10%.

#### 2.4.5 Normalization of BPs

The main problem of using raw bifurcation parameters to study brain dynamics at a single subject level is that, depending on the *G*_*i*_, BPs can have very different ranges (Figure 3.A). Therefore, each set of *BPs*[*G_opt_*] has values that cannot be directly compared between different datasets, irrespective of whether they come from the same or different subjects.

In order to make such BPs comparable, we applied a normalization procedure that i) produced a fixed range of values and ii) generated a “stabilization” of the function 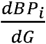, making comparisons between subjects and sessions possible. To this aim we took each *BPs*_*i*_. (set of BPs for a given *G*_*i*_) and split them in two subsets, one containing all the positive values (*pBP_i_*) and another with all negative ones (*nBP*_*i*_). To obtain the normalized bifurcation parameter set (*NBPs*_*i*_) we divided *pBP*_*i*_ and *nBP*_*i*_. by the largest absolute value on that subset. This procedure is given by Equations 10 and 11 and 12.

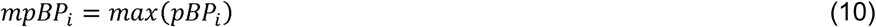

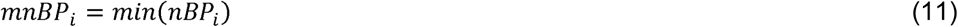

where *mpBP_i_* represents the largest positive value for a given, *G_i_* and *mnBp_i_* the largest negative value for *G*_*i*_. Finally, the normalized vector is given by Equation 12.

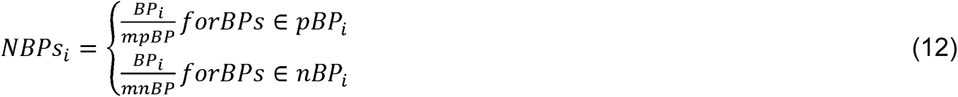

where *BP_i_* represents the raw bifurcation parameter vector and *NBPs_i_* the normalized vector for a given *G_i_*. This simple normalization method yields values between −1 and 1 allowing direct comparison between sessions and subjects.

#### 2.4.6 Types of model-based processing

For this work, we defined two types of analyses:

1. Single recording/subject analysis: In which we calculated BPs using one resting-state scan session/subject.
2. Group analysis: Here we calculated BPs using multiple resting state recordings. The main difference is that in the group analysis the FCM and the frequency spectrum represent the mean of several resting state recordings.

#### 2.4.7 Combinatorial Analysis

An important issue we aimed to address is the minimal quantity of RS information that is required to obtain stable results. To that aim, we implemented a combinatorial analysis in which we increased the number of included resting state recordings. With a complete set of 50 RS recordings the combinatorial analysis consisted of:

1. Doing 50 single recording analyses, one for each recording.
2. Bootstrapping 50 sets of two RS recordings and performing a group analysis on each bootstrap sample.
3. Repeating 2) but for 50 bootstrap samples of 3 RS recordings and increasing the sample size until 20 recordings by steps of 1. Note that this gives a total of 1,000 bootstrap samples (20 combinatorial indexes × 50 analysis).
4. Finally, obtaining the *G*_*opt*_ and its respective NBPs for each of the 1,000 model-based analyses and discarding the ones that did not satisfy our inclusion criteria (KSDist < 0.3 or Fitting > 0.25) for *G*_*opt*_.

## 3. Results

To validate if a dynamic-oriented analysis can be applied to enhance consistency, we first determined if empirical single-subject functional connectivity is consistent through scans. For this, we computed the Pearson correlation coefficient (PCC) for all possible pairs of functional connectivity matrices (CM) both within the same subject and between subjects. In Figure 2 we show that the mean PCC within a single subject (0.67±0.12) is higher than the PCC between participants (0.50±0.10), with an important and substantial overlap of within and between-subjects distributions. While within-subject correlations are on average higher (0.67±0.12) than between-subjects correlations (0.50±0.10), there are many instances in which consistency within the same participant is smaller than the consistency between participants. This mayor issue was tackled from a modeling perspective trying to understand the underlying mechanisms that produce this variability.

**Figure 2:**
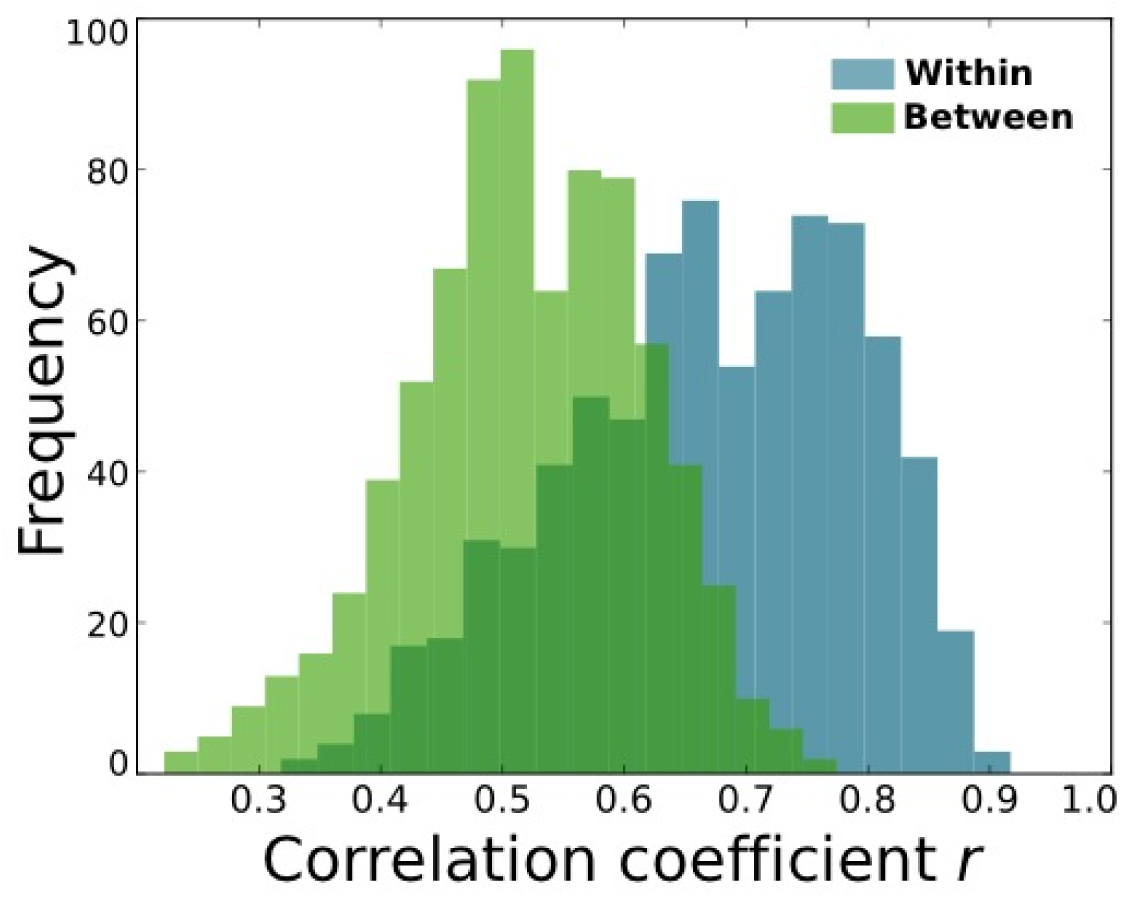
Distribution of Pearson correlation coefficients between pairs of connectivity matrices(CM). Bars in light blue represent correlations for CM of the same subject (within), while in green correlation values for pairs of CM of different participants (between).

We started by exploring the behavior of the intrinsic bifurcation parameters across the extrinsic coupling factor *G*. Intuitively, the collection of parameter values for a given network can be interpreted as a metric that locally represents how brain dynamics are organized. As stated before, a full set of bifurcation parameters (one per region) for a given subject or group can be understood as the Dynamical Core (DynCore). As can be seen from Figure 3.A, raw bifurcation parameters greatly depend on *G*, making it impossible to compare values from different analysis. For this reason, we performed a normalization (see Methods→Normalization of BPs) to obtain DynCores that become invariant for *G* > 2 (Figure 3.B). This allowed the comparison across subjects and sessions in the analysis of brain dynamics. Figure 3.C depicts the change rate of the mean DynCore for 50 different recordings of the same subject. Interestingly, all change rates stabilized for *G* > 2. Hence, if a set of normalized parameters has this degree of consistency, we could then use them to understand local brain dynamics within and between subjects.

**Figure 3:**
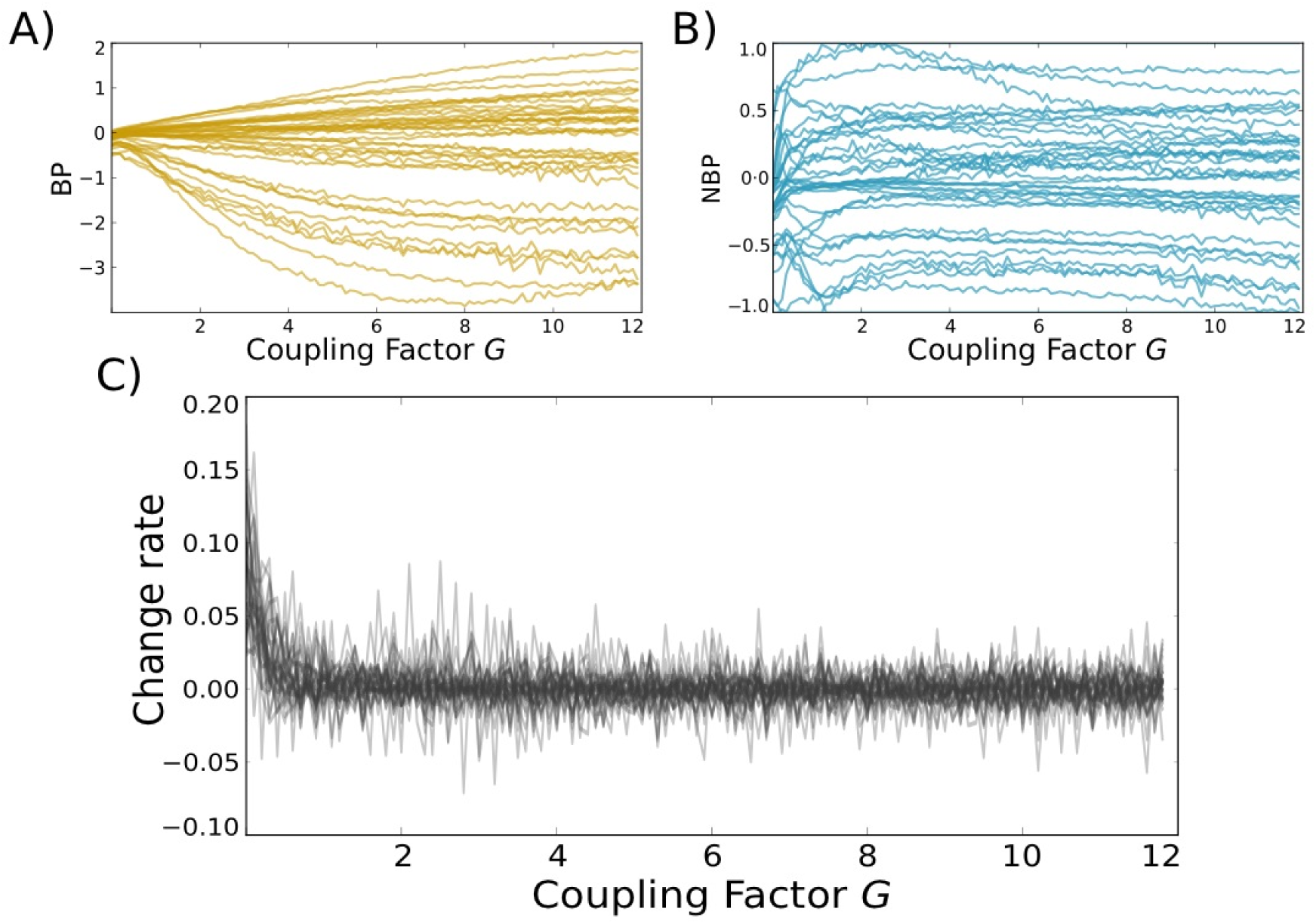
Bifurcation parameter normalization. A) Bifurcation parameter behavior across *G* for asingle subject. Each gold line represents the parameter evolution in a single node. B) The same behavior, but for normalized values (blue lines). Given that normalized values are stable across *G*, they can be used to make comparisons across sessions or participants. C) The mean change rate (derivative with respect to G) for all normalized parameters of 50 different recordings depicted as gray lines, which portray a stable and invariable behavior (slight oscillations around 0) of the average bifurcation parameter across *G* per recording.

To demonstrate that the DynCore is a measure that captures consistent properties of brain dynamics, which can be used to compare both local and global brain dynamics within the same subject, we compared the results of two different within-subject processing pipelines using the same data. In the first one we obtained a DynCore using all 50 RS recordings from the same subject, which is plotted in Figure 4 as a gray-dotted line. We obtained a second set of NBPs by using a combinatorial bootstrap technique that gradually incremented the number of included RS recordings from 1 until 20 (a more detailed explanation in Methods→Combinatorial Analysis), which allowed us to simulate the effect of incrementing scanning time of resting-state information when estimating the DynCore. Boxplots in Figure 4 portray the dispersion of the medians from each of these combinatorial groups. Figure 4 also shows the within subject “mean DynCore” (green lines) obtained by estimating the mean of each box. By looking into local values more closely, frontal regions (among others) exhibited a large diversity of NBPs across all recordings (mean SD: 0.501), whereas temporal and occipital regions showed a smaller parameter dispersion (mean SD: 0.284, 0.277 respectively). Also worth noticing, regions such as the precuneus, parietal cortices, posterior and anterior cingulate gyri as well as the medial frontal cortex showed a mean value near the bifurcation point (0.029 ± 0.016), all hubs with central roles in information trafficking (van den Heuvel and Sporns, 2013).

**Figure 4:**
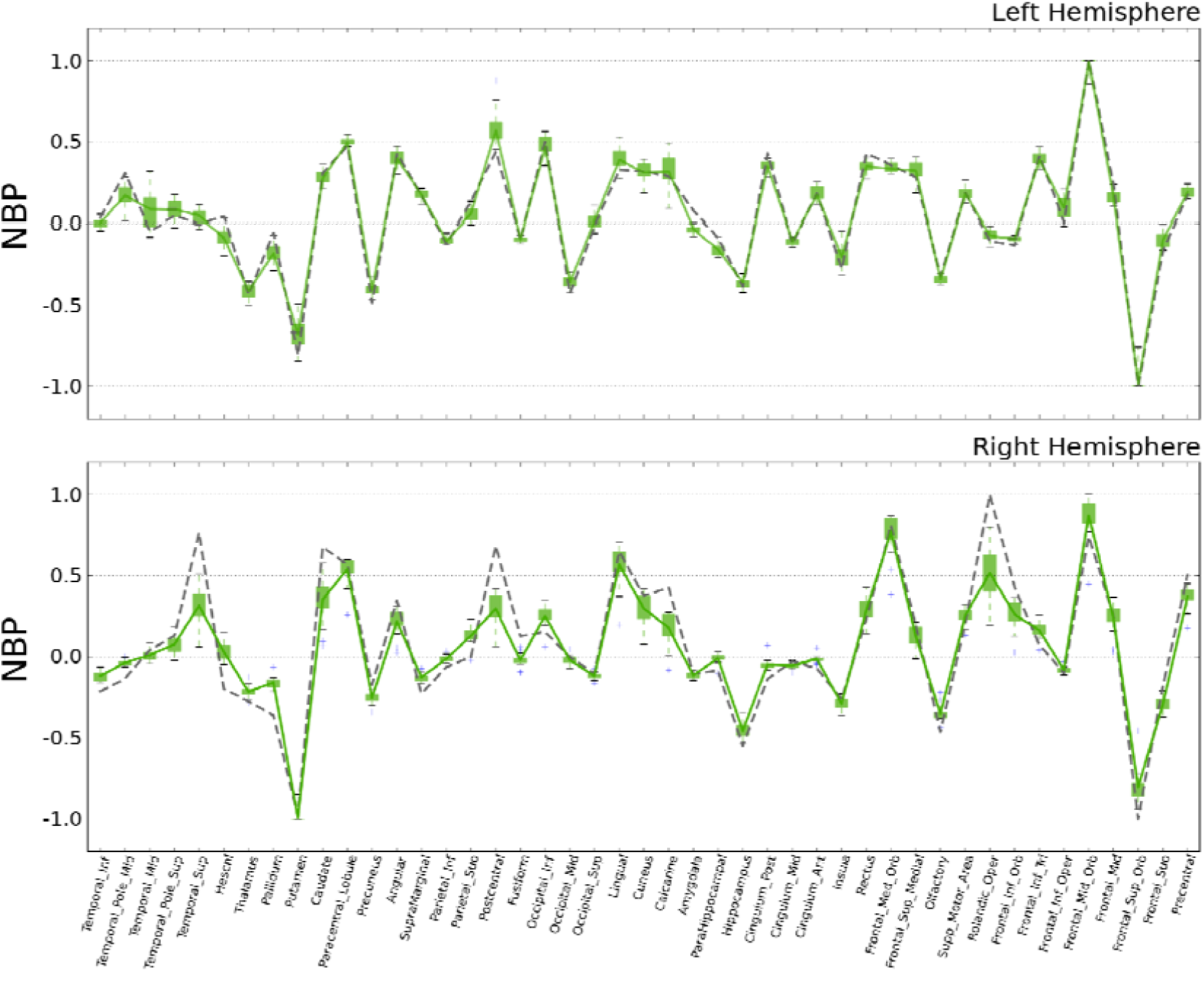
Distribution of the normalized bifurcation parameter (NBP) in each node/region at the (when the simulation best fits the empirical data). Green whisker boxes represent distribution of node medians over all bootstrap samples using data from only one participant. The mean of these random samples (reference DynCore) is represented in green and the gray dotted line is the DynCore (the full set of NBPs) using the within-subject analysis using all 50 RS recordings.

These results also allowed exploring the substantial overlap between the within-subject DynCores obtained by the two methods (r=0.977, p<0.001 (right hemisphere) and r=0.936, p<0.001 (left hemisphere)), suggesting that the DynCore can describe brain dynamics in a consistent way. As the combinatorial analysis yielded estimations as accurate as those computed using all 50 recordings, we decided to use the mean DynCore of the combinatorial analysis as our reference DynCore (green lines, Figure 4) for the later consistency explorations.

To examine the minimum number of recordings that are necessary to reach a reliable estimation, we measured the Euclidean distance from the reference DynCore to median DynCores computed by gradually increasing the number of sessions used for the estimation. This method addresses the impact of increasing information from the same subject on the error rate. Figure 5 shows this evolution for both hemispheres. As more data from the same subject is added to the analysis, the error substantially decreases. Interestingly, this error is quickly minimized and stabilized after adding only four recordings and represents a lower distance than the distance between the reference DynCore and the DynCore calculated from the within-subject analysis using all 50 recordings (gray dotted lines, Figure 4), specially for the right hemisphere, in which the threshold is passed after adding only two recordings.

**Figure 5:**
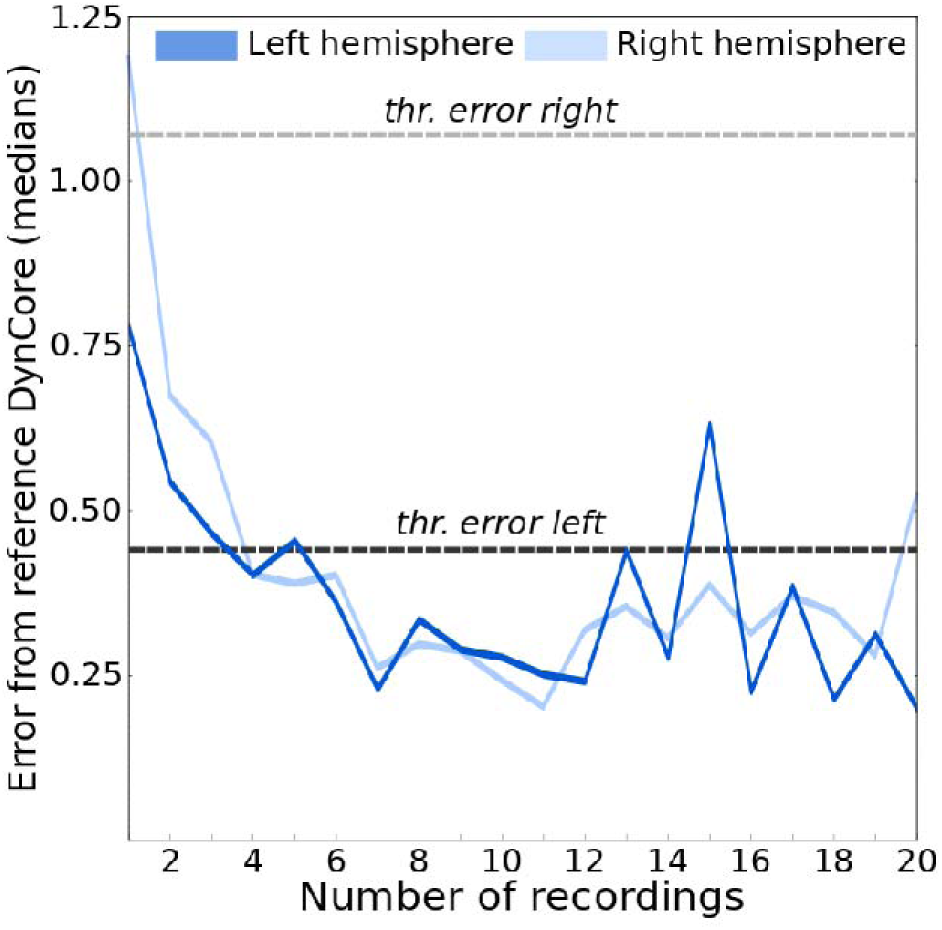
Deviation from reference DynCore as measured by the Euclidean distance. Analysis for the left hemisphere in dark colors and for the right hemisphere in lighter colors. Blue (dark and light) lines represent the distance between the reference DynCore and the median DynCore while increasing the number of recording used to calculate the median. Dotted lines represent the error to reference DynCore from the within-subject analysis using all 50 recordings, here used as a threshold for consistency.

As one of the primary future purposes of this method might be to apply it to individual clinical trials, we studied the error of using raw DynCore estimations (as it could be done analyzing one patient at the time in the clinical environment), without estimating the nodal median first (Figure 4 and 5). We calculated the mean distance of each raw DynCore to the reference DynCore. To show that even raw DynCores (raw bifurcation values) can capture specific and general features of brain dynamics, we also made the exact same analysis using 50 RS recordings from different participants (between-subjects analysis). Similar to what is shown in Figure 5, Figure 6 reveals that the error decreased as more data was added to the analysis both for single subject recordings and also for the between-subjects analysis. Note that although the error never reached levels lower than those estimated using medians (Figure 5), it became relatively low after only four recordings. This is especially true considering that in the within-subject analysis using all 50 recordings (dotted gray lines in Figure 4), the distance was ~0.5 for the left hemisphere and ~1.0 for the right hemisphere (thresholds values in Figure 5), which suggests that using between 15 and 20 minutes of data (3 and 4 recordings) from a single subject, is enough to study local brain dynamics of a single subject in a consistent way (as also suggested by Laumann et al., 2015).

Interestingly, in the left hemisphere within compared to between raw distances required more recordings in order to converge (stars and circles) compared to the right hemisphere, which might indicate a left hemisphere specialization. Another observation of hemisphere specialization is also represented in Figure 5, where the error in the left hemisphere using small number of sessions is smaller than error in the right hemisphere.

**Figure 6:**
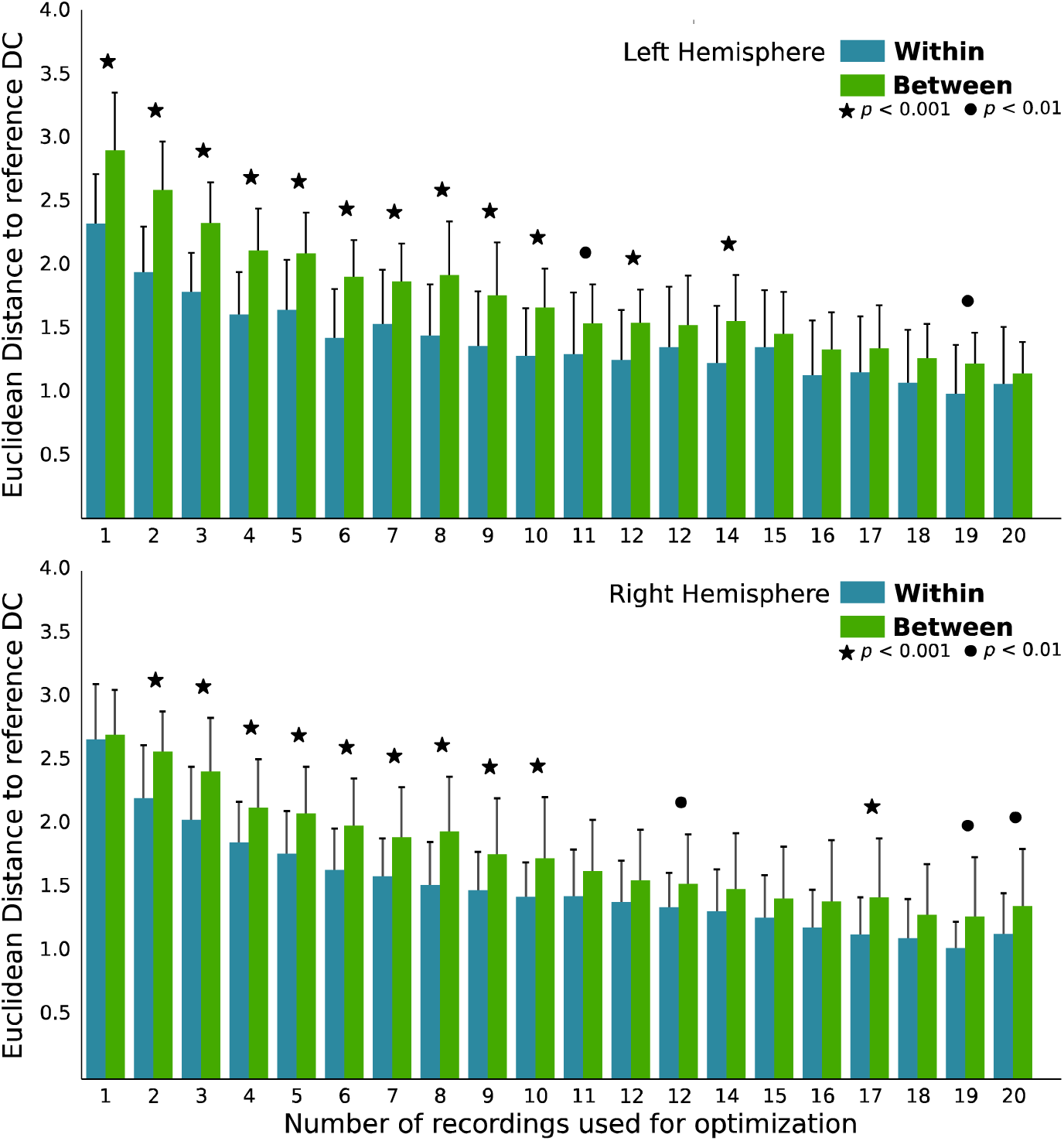
Raw Euclidean distance to reference DynCore for within (light blue) and between (green) subject analysis. This distance represents the mean error to the reference DynCore from the raw normalized bifurcation parameters and plotted as a function of the number of recordings used. Stars and circles represent significant difference values between the within and between-subject Euclidean distances. Finally, using the 1000 raw within-subject DynCores from the combinatorial analysis, we calculated the local error independently for each hemisphere by computing the absolute difference (one dimensional Euclidean distance) to the reference DynCore local value as well as the standard deviation (SD) of all the estimations for each node. As can be seen from Figure 7.A, both estimations yielded very similar results, and as more recordings were added, the estimation became more stable. By computing a correlation between values from the right and the left hemisphere, taking into account all recording points, we found a positive and significant correlation between hemispheres for both measures(*r* = 0.649, *p* < 0.0001 for standard deviation and *r* = 0.562, *p* < 0.0001 for Euclidean distance), indicating symmetry between hemispheres (Figure 7.B). Interestingly and surprisingly, this symmetric profile was damped nonetheless for larger temporal resolutions in the SD case (Figure 7.B, insert). As a general rule though, this analysis suggests that the more recordings are used, the lower the local error measurements are (darker colors in scatter plots).

**Figure 7:**
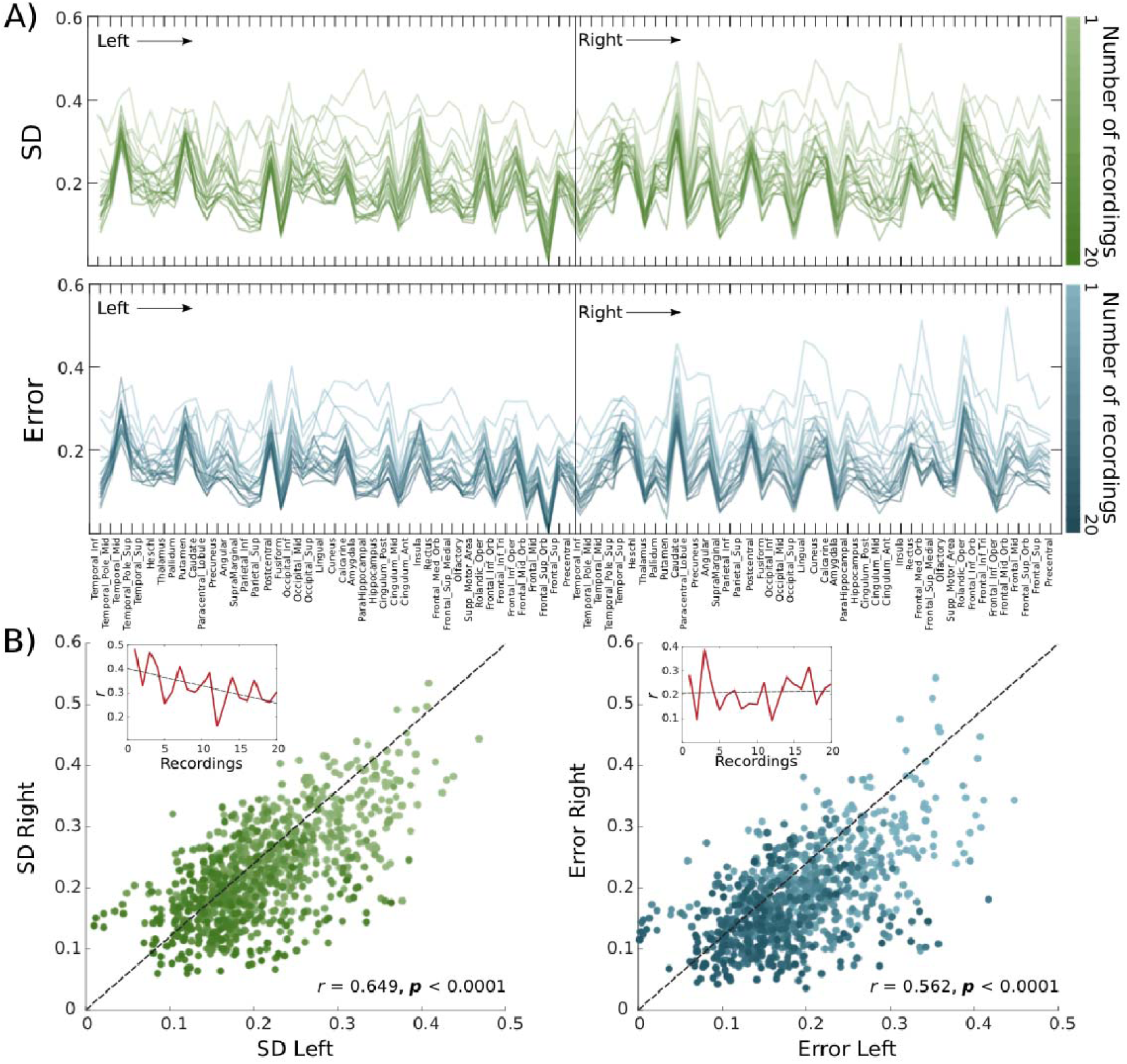
Local error estimation. A) Local standard deviation (green) and Euclidean distance (blue) across number of recordings used for optimization. Color intensity indicates the number of recordings used for optimization (darker is more). B) Scatter plot of local SD and Euclidean distance between the left and the right hemisphere. Correlation value and significance were computed from all local estimations across all number of recordings. Top left inserts show the between-hemispheres correlation value as a function of the number of recordings used, depicting a quantitative estimation for variance and error symmetry.

## 4. Discussion

In this study we used a whole-brain computational model (Deco et al., 2016) to address the consistency of nodal and global metrics of multiple sessions of rs-fMRI from a single subject and single sessions from multiple subjects. We have shown that the collection of bifurcation parameters (the DynCore) reliably reflects brain dynamics of a single subject in resting state, and that these parameters are consistent with group results. Additionally, we found that the estimated error with respect to a group standard as well as local dynamics (nodal errors) converge to their minimum already within four 5-min sessions. We therefore propose that an optimal scan time for analysis of global and local brain dynamics on an individual level is in the order of 15 to 20 minutes. We have also shown that the proposed metrics not only capture the general properties of brain dynamics, but also that DynCores reflect subject-specific features using even only three RS recordings (15 minutes), suggesting an optimal methodological scenario to study brain dynamics at a single-subject level which is important to be able to study neuropsychiatric disorders (Deco and Kringelbach, 2014).

Although some studies have addressed the functional network structure at a single subject level (Power et. al., 2011; Laumann et. al, 2015; Finn et. al., 2015) and it is well known that the brain presents a high level of variability in terms of cognitive domains (Anderson et. al., 2013), resting state functional connectivity profiles (Biswal et. al., 2010; Mueller, 2013) and structural connectivity (Bürgel et. al., 2006), most of the information generated so far about resting state in disease comes from group analysis (see Zhang and Raichle, 2010) in which subject-specific features might be ignored by averaging out local and subtle differences. Further, single-subject resting state dynamics can be used as fingerprints for subject identification (Finn et al., 2015), which stresses the importance and consistency of single-subject data analysis.

Our results suggest that there is a trade-off between data consistency and scanning time, though highly consistent results can be achieved with reasonably short scanning times of about 15 minutes. This is not the first study trying to address reproducibility and consistency in resting state sessions. Many studies have shown that the longer the scanning time is, the better the consistency and reliability of the analysis is. Interestingly, some have found that the optimal scanning time to have reproducible results is greater than 14 to 20 minutes (Birn et. al., 2013; Anderson et. al., 2011; Laumann et al., 2015), which represents around three to four sessions in the present study, corroborating the idea of a minimal scanning time to have reproducible results. To extend this, another study (Kalcher et al, 2012) found that as the scanning time increases, the number of independent components found also increases monotonically also suggesting a better estimation of resting state dynamics by longer scanning times.

As within-subject data in this study was acquired in different days in a period of six months, the estimation of consistency could lead to an over-estimation of the optimal minimum single-session scanning time as day-to-day variability might play an important role. In a clinical environment though, it seems more feasible to acquire 15-20 minutes in a single session than to split it up in different scanning sessions. Moreover, this split across different days also means that the day-to-day variability might not only produce an over-estimation, but also produce valuable information about changes in brain organization only captured over longer periods. Future works should then analyze the consistency between long-length single session DynCores and multiples-sessions DynCores.

Because the reliability of brain network metrics derived from resting state is highly dependent of both the initial parameters and the network density (Braun et. al., 2012), the current study suggests a novel and rather powerful approach that not only globally but locally (Figure 7.A) reduces the estimation error, opening new possibilities for a systematic regional analysis across participants. Our results are in line with many findings addressing these issues (Kalcher et al 2012; Birn et. al., 2013; Anderson et. al., 2011) and in contrast to what now has become a common practice, five minutes (van Dijk et. al., 2010) of scanning time may not be sufficient to produce reliable results on an individual level as in fact, the reduction of error in the estimation of local bifurcation parameters in the present study could also be explained by the fact that the longer the scanning time, the higher the signal-to-noise-ratio (Murphy et. al., 2007).

Taking this into account, the need to study individual patterns in clinical trials becomes more relevant. Extending our findings that the exploration of brain dynamics at rest can be performed at the subject level, the same latter study found that these functional fingerprints are unique and are bounded to subject’s intrinsic connectivity properties (Finn et. al., 2015). Because the method proposed here also allows identifying consistent individual changes in local dynamics with a relatively low number of sessions (see Results), follow up analyses should study disease by means of the DynCore, focusing on local and global changes that might be subject-specific, hence giving a first proxy for a potential sensitive biomarker. Also interesting is the fact that although we found symmetric profiles both for local standard deviation and error, symmetry for standard deviation was better explained by fewer recordings and weakened by adding more (Figure 7.B). This suggests that although using multiple sessions is better to obtain a highly consistent estimation of the underlying functional organization of brain at rest, a plausible very slow physiological and/or environmental mediated variability (Taubert et al., 2011) of brain states could be overlooked using this approach.

To explore this idea further, it is worth remarking that the correlation value between hemispheres has a tendency to go down as more recordings are added in the standard deviation (SD) estimation (Figure 7.B). This might suggests that nodal variability might be explained by two main components. First, a physiological slow-oscillatory component expected and second Gaussian noise component caused by technical and/or technological limitations. It is possible that, when a small number of sessions are being used, the physiological component is larger compared to the Gaussian noise and hence a symmetric profile is observed. As more sessions are added, the Gaussian component becomes much larger, which in turn causes a reduction in the observed symmetry. This might indicate that to study very slow changes in the brain’s functional organization, longitudinal RS sessions must be processed in series of small batches. Like this, symmetry in variability patterns (error to grand-average DynCore and SDs) can reveal useful information to study changes in the brain at rest that occur over long periods of time (Taubert et al., 2011) that could ultimately be related to different, and not well studied physiological, cognitive, emotional or even pharmacological factors.

At a whole brain scale, the median value of the reference DynCore is 0.017, which indicates a critical oscillatory behavior optimally operating at the edge of a bifurcation (Deco & Jirsa, 2012; Deco et al., 2016). By exploring bifurcation parameter values node by node, we were also able to identify local variability. Frontal regions seemed to exhibit larger parameter dispersion (larger standard deviation) compared to occipital and temporal cortices, suggesting that the functional variability of frontal regions might be larger compared to other parts of the brain. Interestingly, a study (Anderson et a., 2013) found that functional diversity in terms of cognitive domains varies across the brain. More importantly, frontal regions presented higher levels of diversity compared to other regions such as temporal and parietal cortices. A more specific and causal interpretation of local bifurcation values is beyond the scope of this study, but it is worth noticing that at least qualitatively, local NBPs display patterns that seem to relate with functional segregation. Also supporting this idea is the fact that after performing a simple *post-hoc* analysis, nodes from the default mode network (as in van den Heuvel & Sporns 2012, 2013) display a mean parameter value of 0.02, which is significantly closer to zero (*p* = 0.041) compared with the same mean value of 10,000 randomly bootstrapped subnetworks of the same size. Given that all these regions are considered hubs (van den Heuvel & Sporns 2012, 2013; Tomasi & Volkow 2010), this suggests that local values closer to a transition or bifurcation state might also have central key role in information transfer. Future studies should explore with greater detail the functional role of local bifurcation dynamics in information trafficking and cognition.

A primary limitation of the present study is the lack of single-subject structural connectivity matrices. Because current models (including the one presented here) of brain dynamics are based on the average structural information (Deco et. al., 2013), it would be interesting to investigate if the number of sessions required to optimally reach the reference DynCore is smaller by using single-subject structural matrices. Finally, due to technical limitations, the temporal resolution is relatively low. Further modeling of resting state activity using MEG (Nakagawa et al 2014) at a single subject level should also give richer insights into the behavior of local dynamics both in health and disease.

## 5. Conclusions

We presented a conceptual framework for analyzing local brain dynamics at a single subject level by modeling resting state activity with a supercritical Hopf bifurcation model. The estimated intrinsic parameter, the DynCore, allowed us to explore the amount of information required to minimize the error in order to have consistent results reflected by the Euclidean distance evolution across sessions added. Overall, including four sessions for the same subject yielded highly consistent results. Additionally, we showed that adding more recordings unexpectedly reduced hemispheric symmetry of error estimations of local dynamics. These methods open new avenues to analyze local brain dynamics at a single subject level that may reveal localized and subtle changes in health and disease.

